# Keratin5-BMP4 mechanosignaling regulates reciprocal acetylation and methylation at H3K9 to define blastema cell remodeling during zebrafish heart regeneration

**DOI:** 10.1101/2020.09.10.290882

**Authors:** Xuelong Wang, Huiping Guo, Feifei Yu, Wei Zheng, Hui Zhang, Ying Peng, Chenghui Wang, Gang Wei, Jizhou Yan

## Abstract

Heart regeneration after myocardial infarction remains challenging due to scar and ischemia-reperfusion injury. Here, we show that zebrafish blastema regeneration can effectively resalvage the wound myocardium and blood clot through cytoplasmic exocytosis and nuclear reorganization. The cell remodeling process are also visualized by spatiotemporal expression of three core blastema genes: alpha-SMA- which marks for fibrogenesis, Flk1for angiogenesis/hematopoiesis, Pax3a for remusculogensis, and by characteristic chromatin depositions of H3K9Ac/H3K9Me3. Genome-wide enhancer identification links the depositions of the two histone marks to the chromatin state and these three core blastema cell phenotypes. When the blastema subcellular fractions are introduced into the cultured zebrafish embryonic fibroblasts the altered transcription profile is comparable to the blastema transcription in terms of extracellular matrix structural constituent, vasculature development/angiogenesis, and cardiac muscle regeneration. From the subcellular fractions we identify 15 extracellular components and intermediate filaments, and show that introduction of human Krt5 and noggin peptides conversely regulate PAC2 cells F-actin reorganization, chromatin depositions of H3K9Ac/ H3K9Me3 and phosphorylation of Smad, which are accompanied by characteristic transcriptions of *bmp, bmp4*, three core blastema genes as well as specific histone acetylation/methylation-related genes. Collectively, this study establishes a new Krt5-BMP4 mechanosignaling cascade that links extracellular molecules to chromatin modifications and regulates blastema cell remodeling, thus providing mechanistic insights into the mesoderm-derived blastema regeneration and underlining a therapy strategy for myocardial infarction.

## Introduction

Coronary artery disease (CAD) also known as ischemic heart disease is the most common type of cardiovascular disease. Limitation of blood flow to the heart brings about cell oxygen starvation (ischemia), and cardiomyocyte necrosis (myocardial infarction, MI). The death of cells then triggers a cascade of local inflammatory response and fibrogenesis (Bing, 2001; Smit et al., 2019). If the impaired blood flow is not deteriorated, myofibroblasts and endothelial cells proliferate and migrate into the infarct zone, where myofibroblasts deposit a network of collagen to form a hypocellular collagenous scar (Krijnen et al., 2002; Virag and Murry, 2003). The resultant myocardial scarring could impede conduction velocity and cause lethal arrhythmias. Besides scaring, another common problem is the second round of ischemia-reperfusion injury (IRI) or reoxygenation injury after a period of ischemia or lack of oxygen and nutrients from blood (Parviz et al., 2019). Therefore optimal myocardial infarction therapy requires a scarless regeneration, which can balance inflammatory repairs, blood resupply and myocardial renewal.

Gene edition of periostin-expressing myofibroblasts could reduce collagen production and scar formation (Kanisicak et al., 2016), and forced viral-mediated gene expressions or chemical cocktails could induce reprograming of mouse non-muscle cells into cardiomyocytes (Fu et al., 2015; Xin et al., 2013). More inspiringly inflammatory cytokines, such as interleukin-1beta (IL-1beta) and tumor necrosis factor-alpha (TNF-alpha), could induce the transformation of human dermal microvascular endothelial cells into myofibroblasts in skin fibrogenesis (Chaudhuri et al., 2007; Karasek, 2007), and the (myo)fibroblast phenotype could reverse back to the precursor phenotype (LeBleu et al., 2013; Willis et al., 2006). These data elicit an anticipation to convert the inflammatory reaction and inflammatory cells into cardiac tissue regeneration (Zlatanova et al., 2016).

Zebrafish can faithfully repair the complicated tissue organs through blastema regeneration (Govindan and Iovine, 2015; Lepilina et al., 2006; Wang et al., 2012). Several lines of evidence indicate interweaves between blastema formation and myofibroblast formation. Historically, both myofibroblast and blastemal cells are characterized by fibroblast phenotype with increase of extracellular matrix. They are heterogeneous in the cell origin and plastic in the phenotype. Myofibroblasts can arise from conversion of local resident fibroblasts, differentiation of bone marrow-derived progenitors, endothelial-mesenchymal transition (EndMT) and epithelial-mesenchymal transition (EMT) (Bochaton-Piallat et al., 2016; LeBleu et al., 2013). Likewise heterogeneous blastema cells in lower jaw regeneration originate from fibroblast cells, nucleated blood cells, fragmented muscle cells, pigment cells, and local vascular system (Wang et al., 2012; Zhang et al., 2015). Different from fibrogenesis with accumulation of scar tissue, blastema proceeds with tissue respecification. Thus, zebrafish blastemal regeneration strategy could provide a clue to switch inflammatory repair to cell plastic regeneration.

Here, we employed transgenic reporter fish and cell identity analysis, traced the fate of the injured myocardial cells and blood clot, and investigated molecular mechanisms responsible for heart blastema regeneration in zebrafish. Our findings at subcellular and molecular levels revealed dual regulation of cytoplasmic intermediate filament mechanosignaling and histone modification switch on cell cytoplasmic remodeling and nuclear reorganization pertain to heart blastemal regeneration.

## Results

### 1. Epicardium and subepicardium derived cells invaded and induced fibrinolysis and blastema formation

The previous studies showed that zebrafish can vigorously regenerate the amputated cardiac muscle and restrict scar formation (Lepilina et al., 2006; Poss et al., 2002). However, these studies did not clarify how zebrafish renovate the thrombus and reconstruct the blood supply system. In this study, we found that resection of the ventricular apex induced bleeding, blood clot formation, and wound healing. As proliferation and migration of pericardial cells sealed the wound and the blood clot, the resultant pericardial-covered clot mix (hereafter called epi-clot) established a pre-blastema field, which grossly contoured the resected apex in morphology (Fig.1, Fig.1). From 2 to 3 days post-amputation (dpa), the adjacent subcardial cells invaded into the epi-clot, and triggered tissue lyses, fibrogenesis, and blastema formation. Concurrently with myofibroblast formation, blastema formation but progressed with vascuologenesis and cell remodeling. Meanwhile new types of light staining cells (or pre-blastema cells) emerged from the lysed tissues and clot, and then transformed to heterogeneous blastemal cells. By 8dpa, the pre-blastema field was filled with a variety of blastemal cells as well as the residual epi-clot and hypocellular mus-clot. During blastema reformation/respecfification from 10dpa to 30dpa when the myofibrills remuscularized the blastema and encapsulated the residual clot, the inside nucleated blood cells (nbc) were induced and converted into various types of hematopoietic-derived cells in morphology. Here we not only confirmed the previous observations that epicardial and subepicardial cells were actively involved in blastema formation (Poss et al., 2002), but also displayed the cell remodeling processes through integration of fibrinolysis, musculolysis, fibrogenesis, revascularization/ hematopoiesis, and remuscularization.

**Figure 1.**
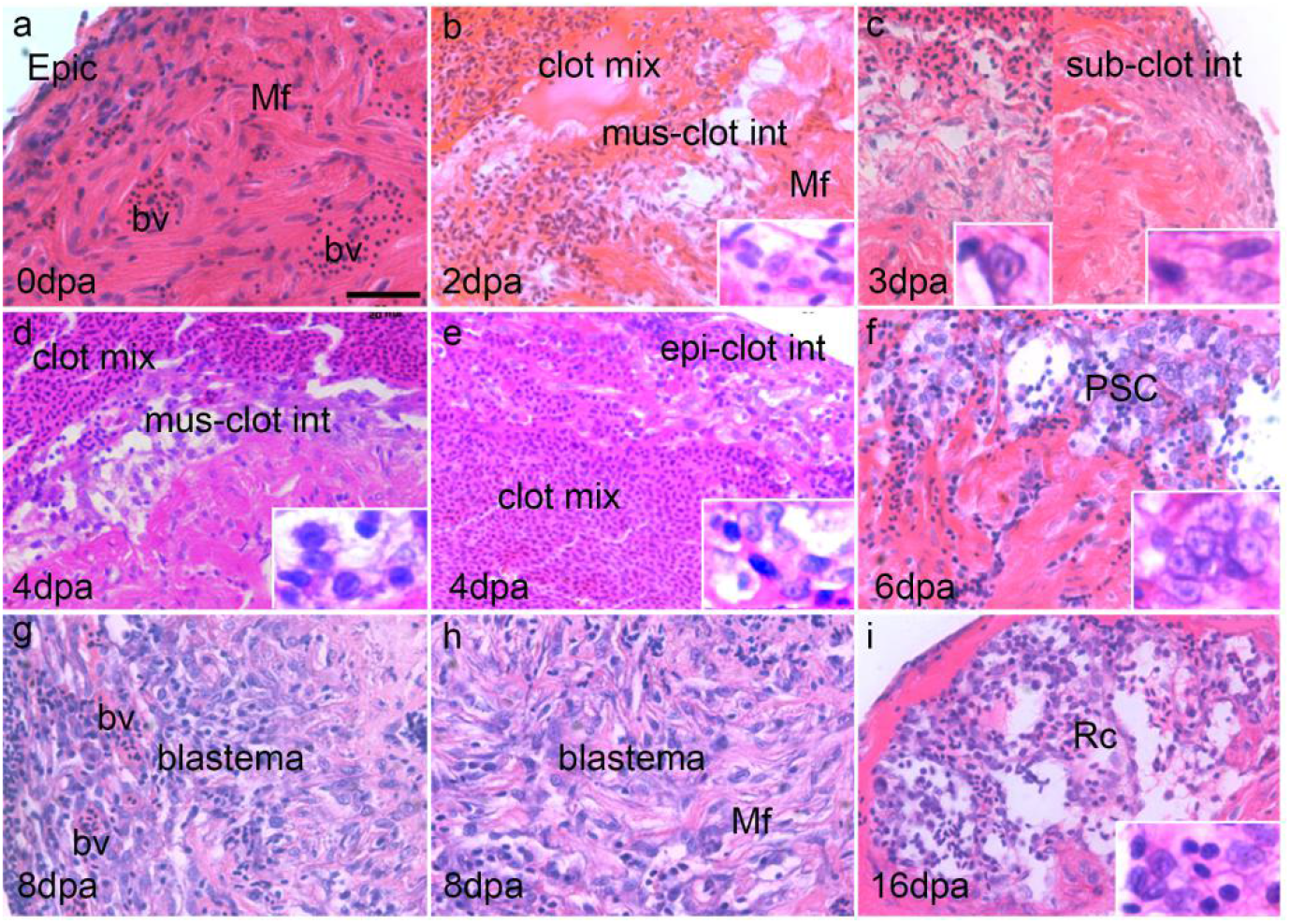
Histological observations on blastema formation and reformation during heart regeneration After amputation of the ventrical tip, an initial response was bleeding and hemostasis. In the following days the epicardial cells immigrated and incorporated with the outmost fibrined nbcs to form a pericardium-like outer layer covering the nbc clot. Tissue lyses began along the intersection between epi-clot (epi-clot int), subcardium (sub-clot int), and the damaged muscle (mus-clot int).The light cells refer to those cells that showed hematoxylin-resistant and had a large nucleus with vacuolar cytoplasm. As more subepicardial cells invaded, the light-staining areas progressively expanded toward the epi-clot and the mus-clot. Subsequently the light cells became basophilic and stem cell-like in morphology. With increase of blood vessels, a typical blastema was formed by mass of basophilic and fibroblast-like cells with excess of extracellular matrix. As shown in the islets, there were two size types of light cells. The large light cells had abundant cytoplasm and a variety of vacuoles; the small light cell contained small amount of cytoplasm and a relatively *large* intact nucleus (N). Bar = 200 μm.

### 2. Blastema cells undertook cytoplasmic exocytosis and nuclear reorganization

Compared to the histological sections the transformation processes of blastemal cells were more discernible at the subcellular level under transmission electron microscopy (Fig.2). The blastemal cells arisen from the myocardial cells underwent cytoplasmic vacuolation and efflux, and reproduction of new cytoplasmic components. In contrast to the extensive cytoplasmic autolysis, the nuclei displayed resistance to degradation. Such nuclear reorganization were more obvious within the activated nbcs, initiated from nuclear condensation and heterochromatin formation. Noticeably certain induced blood clot cells exhibited the morphological characteristics similar to the presumptive hematopoietic stem cells (PSCs) described by Rubinstein (Rubinstein and Trobaugh, 1973). Thus, our histological and ultrastructural observations clearly showed the blastema cell resalvage processes from the cardiomycytes and clotted nbcs through cytoplasmic exocytosis and nuclear reorganization during blastema regeneration.

**Figure 2.**
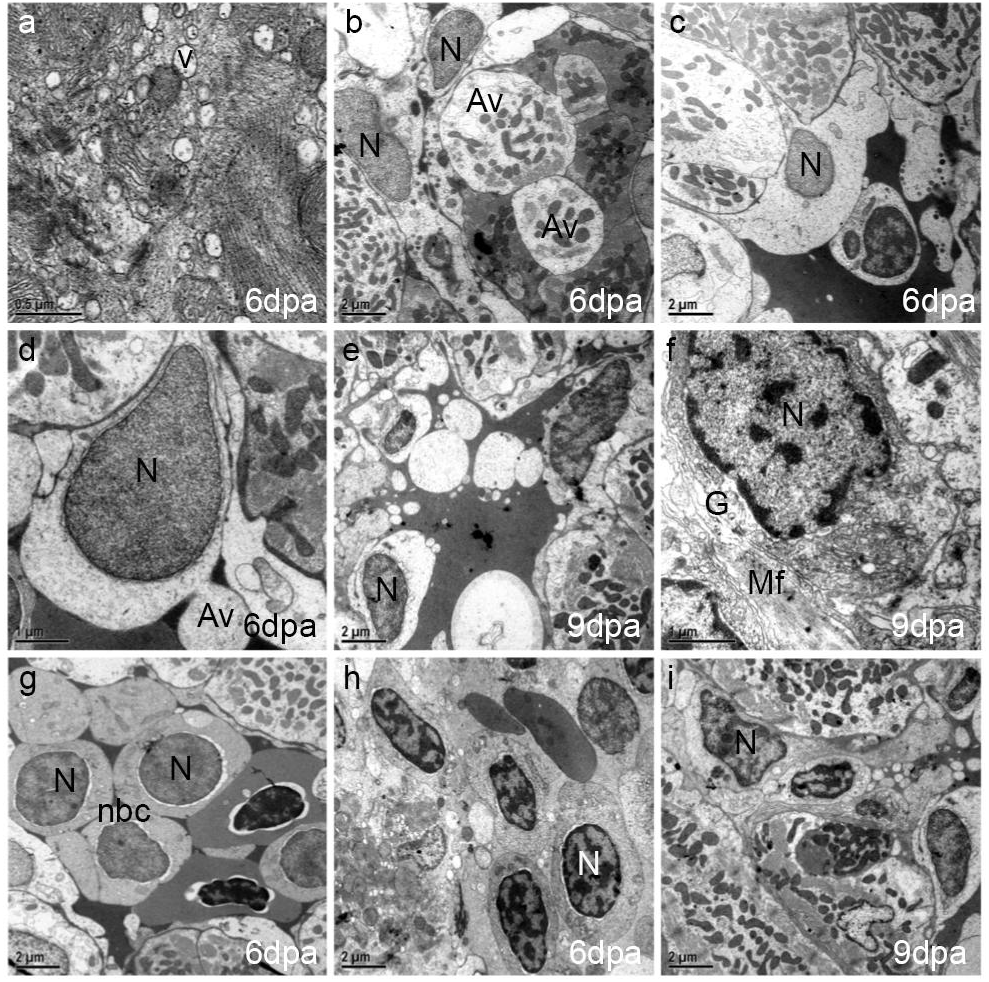
Ultrastructural observations on the cytoplasm remodeling and nuclear reorganization of blastema cells a-f, The large light cells contained numerous transparent vesicles. These membrane-bound vesicles were limited by a double or single membrane and usually filled with granules or other inclusions. Some vesicles were fused into bigger vacuoles, or eliminated from the cells. The chromatin lining the nuclear periphery throughout the nucleus became condensed. The nuclear pore-like structure became evident due to complicated folding of the nuclear envelop. Endoplasmic reticula and Golgi apparatus aggregated around the poles of the nucleus. Mitochondria and myofibrils were scattered throughout the myocyte organnells. g-i, Within the nucleated blood cells (nbc), the nucleus was initially condensed, and left an empty rim around the nucleus. Then the nucleus was enlarged with increasing thick clumps of chromatin beneath the nuclear membrane. (N) nucleus, (No) nucleolus; (M) mitochondria; (Mf) myofibrils; (G) glycogen; (v) vesicles, (Av) autophagesome-like vacuole. Bar: a=0.5μm, d=1μm, the rest = 2 μm

### 3. Three tissue specific factors tracked myofibroblast and blastema formation

According to the previous report, zebrafish adult cardiac muscle regeneration is comparable to embryo cardiac morphogenesis from primitive embryonic structure into three mature myocardial lineages of primordial, trabecular and cortical cardiomyocytes (Gupta and Poss, 2012). To trace the origin and phenotypic respecification processes of blastema cells, we utilized isl1:GFP and flk1:GFP transgenic fish to visualize isl1^+^- and flk1^+^- GFP fluorescence in the injury site; and also used antibody staining for myofibroblast marker-alpha-smooth muscle actin (alpha-SMA) and Pax3a protein to visualize the proliferating myofibroblasts and myogenic progenitors, respectively. Despitesignal of isl1-GFP undetectable, the spatiotemporal expression patterns of alpha-SMA, Flk1 and Pax3a were roughly paralleled to blastemal fibrogenesis, angiogenesis and myogenesis, respectively (Fig.3A). These results indicated that the three genes could represent and mark the blastema formation and tissue respecification processes.

**Figure 3.**
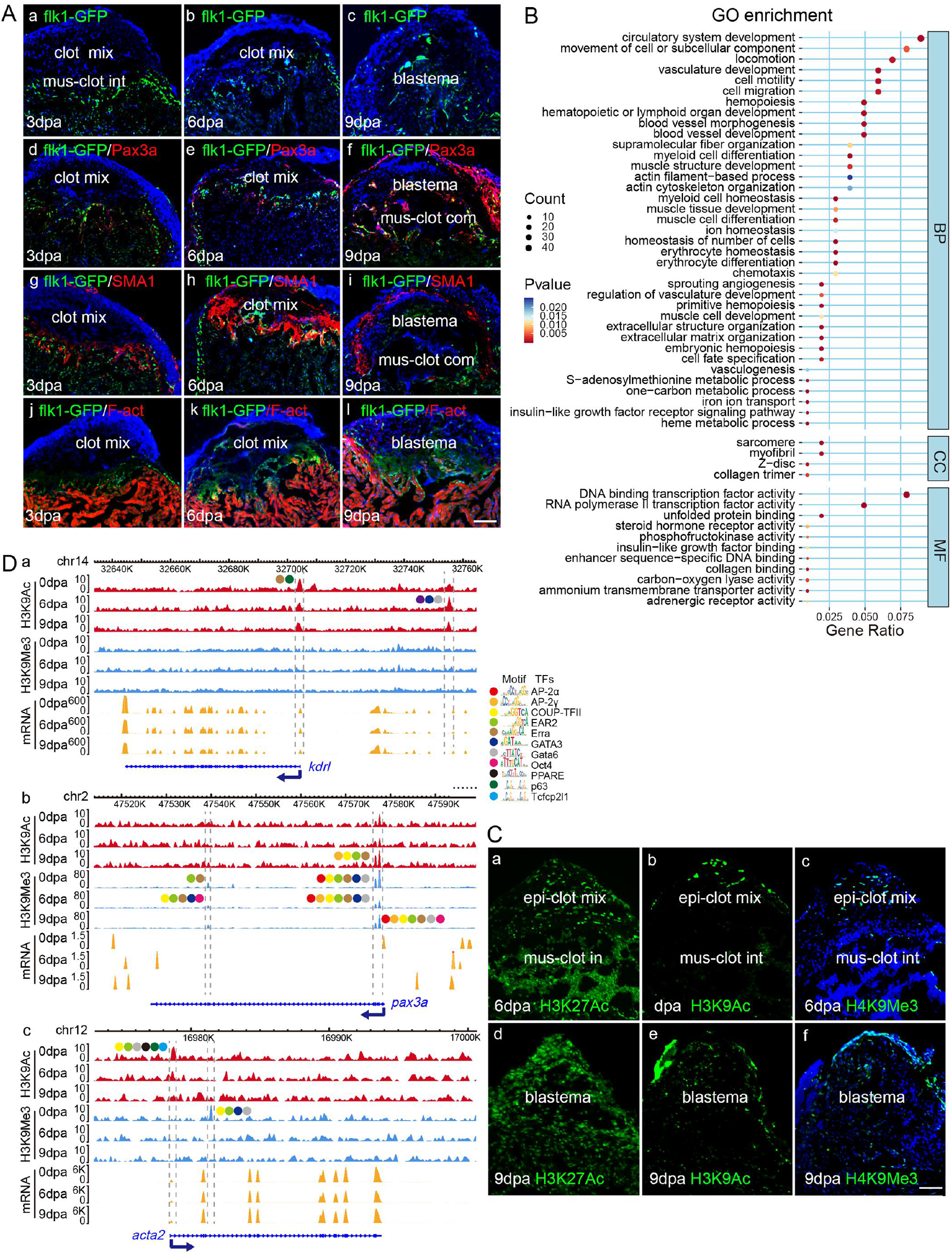
Blastema phenotype and genotype and regulation at chromatin level A showed immunohistochemistry (IHC) tracking of blastema cell origin and respecification. At 3dpa only a few of Flk1-GFP cells and a-SMA-expressing cells emerged from the damaged muscle and the subepicardium, then progressively penetrated to blastemal center and the fibrin clot, and expanded to the whole precardial field. By 9dpa, alpha-SMA expression was only detected in the outskirt of the blastema, presumptively in regenerating subepicardium. Pax3a expression was consistently increased in the regenerating epicardium and outskirt of the blastema, paralleling with epitheliazation and masculaturation in the regenerate. Pholiding-labeled F-actin indicated cardial muscle staining. B showed the top enriched BP terms by GO enrichment analyses of 626 significantly differential expression genes. C showedrepresentative photography of H3K9Ac and H3K9ME3 immunohistochemistry. Anti-H3K27Ac antibodies were extensively stained in the whole cells. In contrast, H3K9Ac and H3K9ME3 were spatiotemporally stained in the regenerating cells. At 6dpa, intense H3K9Ac signals mainly aggregated at the outmost epicardium and less in muscle-clot mix in contrast to the enhanced H3K9Me3 expression in the epi-clot mix. At 9dpa, both H3K9Ac and H3K9Me3 expressing cells increased but differentially distributed over epi-clot mix (blastema region). D showed H3K9Ac/Me3-specific enhancers and enhancer motifs at three core blastema genes. Dot lines indicate the enriched peaks. Color balls indicate sites of enhancer motifs (blastema respecification regulators). Bar = 50μm

To unveil the genotype-phenotype correlation during the blastema cell remodeling process, we conducted time-point based RNA-seq analyses and identified 626 significant transcripts between three time points(P<0.05). Functionally these genes comprehensively reflected the blastema regenerative progress as above histomorphological descriptions (Fig.3B). Although transcriptions of *kdrl* (*flk1*), *acta2*, and *pax3a* were not included in the 626 significant transcripts, their equivalent functions were fully exhibited by the category of gene ontology of the 626 genes (Fig.3B) in contribution to the extracellular matrix organization, angiogenesis/ hematopoiesis, and muscle structure development (Table S1). Therefore we concluded that *kdrl, acta2*, and *pax3a* represent three core blastemal tissue specific genes in regulation of the blastemal cell remodeling process.

### 4. H3K9Me3 and H3K9Ac switch dynamically marked the blastema cell proliferation and tissue respecification

Considering epigenetic regulation as a bridge between genotype and phenotype, we examined the distributions of 8 common epigenetic marks during heart regeneration using immunohistochemistry (Fig. S2). Among the tested histone modification marks H3K9Ac and H3K9Me3 exceptionally exhibited a dynamic change in the immunostaining of cells from the epicardium to the blastemal center. Intense H3K9Ac territories along the epicardium showed less staining for H3K9Me3, while dense H3K9Me3-staining foci within the blastema center showed week H3K9Ac signals (Fig.3C). Subnuclear localizations showed an intense punctual staining of H3K9Ac in the nuclei of mus-clot cells, in contrast to intense H3K9ME3 signal in DAPI-heavy pericentric heterochromatin of the clotted nbcs (Fig. S3). We gathered that the spatiotemporal staining pattern of H3K9Ac was coincidence with the cell proliferation route along the epicardial cells to the blastema cells while H3K9Me3 staining patterns accompanied blastema respecification from blastema center myofibroblast/blastema formation to blastema outskirt remuscularization. When the regenerating heart was pretreated with H3K9Me3-specific and H3K9Ac antagonists, either H3K9Me3 or H3K9Ac reduction impaired blastema formation as indicated by the Flk1-GFP and DAPI staining signals (Fig. S4). These results suggest that H3K9Me3 and H3K9Ac coordinately regulate blastema cell proliferation and phenotypic respecification.

### 5. Genome-wide identification of potential regulatory sequences involved in chromatin state dynamics

The patterns of H3K27Ac and H3K27Me3 profiles along the genome were previously used to identify potential enhancers and promoters in particular cells (Hawkins et al., 2011). By the same way, we performed genome-wide ChIP-seq analyses to identify the potential regulatory elements of the H3K9Ac/Me3 marks. The distribution of the two chromatin-predicted enhancers was primarily distal to the TSSs, with totally 82.84% lying in intergenic regions, and 3.69% falling in at promoter-TSS sites. The intergenic regions enriched 28.03% of H3K9Ac and 75.39% of H3K9Me3, compared with 0.19% of the H3K9Me3 and 15.06% of H3K9Ac poised at promoter-TSS sites (Fig. S5).

Consistent with the cell-specific and subnuclear staining patterns, the two histone marks were mutually exclusive and coordinated at the intergenic and promoter-TSS sites, including at the core blastema genes (Table S2, Fig. 3D). H3K9Me3-dominant sites showed less enrichment for H3K9Ac while those dominantly marked by H3K9Ac were less enriched for H3K9Me3 (Fig. 3D, Fig. S6). The reciprocal situation was also confirmed by ChIP-PCR evaluation of the enrichment of the two histone modification marks at the promoter regions of several interest genes, particularly obvious at the *pax3a* and *acta2* genes (Fig. S7).

The majority of enrichments at intergenic regions and multiple binding sites at the nearest genes conformed to the structural binding mode through those persistently enriched constitutive enhancers and temporally enriched enhancers. Relatively those moderately H3K9Me3 marked genes were more temporally variable than the highly H3K9Ac-marked genes during the three time-points of regeneration (Table S3). Statistical analyses of the top 1000 of the enriched peaks demonstrated reverse correlation between the two histone mark depositions (r=-0.56) and multifaceted relationship between the H3K9Ac/Me3 depositions and the targeted gene transcriptions. Only a part of targeted genes’ transcription exhibited positive correlation with H3K9Ac enrichment, or negative correlation with H3K9Me3 enrichment (Fig. 3D, Fig. S6, Table S3).

### 6. H3K9ME3 and H3K9Ac-poised enhancers correlated with the key blastema regulators

To further analyze how H3K9Ac and H3K9Me3 enriched enhancers regulate the targeted genes, we investigated if known transcription factor binding sites (TFBS) from the JASPAR and TRANSFAC databases were enriched at predicted enhancers in a histone modification-specific manner (Fig.S8). The high-confidence (p< 1E-50) motifs were constantly found in both datasets and remained stable at the three time points. These constant motifs included two marks-shared (p73, p63, p53, Tcfcp2l1(CP2), Coup-TFII, Ear2, Prdm1, Erra), H3K9Ac-dominant (AP2, FXR), and H3K9Me3-dominant (CTCF and Gata6). In addition, Table S4 listed the high percentage of Target Sequences (>10%) and temporal-sensitive motifs, such as Cdx2, Cdx4, Foxa2, FoxL2, Gata3, Gata4, HIF-1b(HLH), Hoxa11, Hoxb13, Smad3(MAD), OCT:OCT-short (POU, and Homeobox). Of these transcription factor motifs that we classify at H3K9Ac/Me3-specific enhancers, almost all of the corresponding factors are known to play a role in the regulation of cellular differentiation and organogenesis during vertebrate development, particularly in myocardial differentiation (Gata4, Gata6), endothelial cell biology (Gata3), hematopoiesis (Cdx-Hox pathway) (Kikuchi et al., 2010; Zhou et al., 2008). We designated these H3K9Ac/Me3-specific enhancer motifs as blastema respecification regulators.

As AP-1 has been shown to regulate chromatin accessibility changes and gene expression programs during cardiomyocyte regeneration (Beisaw et al., 2020), most of the blastema respecfiication regulators were mapped around three core blastema genes in a spatiotemporal manner(Fig. 3D, Table S5). Of the top 20 enriched TF-motifs, the H3K9Ac-enriched TFs (p63, Erra, Tcfc2p21, Gata3, Gata3, Foxk1) mapped to gene *kdrl*, Ear2, Erra, Coup-TF11, AP2, Foxo3, Oct4, Gata6, Gata3 mapped to *pax3a* in a H3K9me3-dominant manner, while gene *acta2* was only targeted at 0dpa by two marks-enriched TFs (Tcpcp211, p63, Ear2, Coup-TF11, Gata6, Gata3). Significantly these dynamic positioning of histone marks-specific enhancer motifs at the three core blastema genes conformed to their spatiotemporal expression patterns shown in Fig. 3A and Fig.3C, i.e. progressive activation of *kdrl*, transient activation of *acta2* and multivalent *pax3a* activations.

### 7. Identification of key effectors responsible for blastema cell nuclear reorganization and cytoplasmic remodeling

Since cell-derived extracellular vesicles (EVs) are powerful conveyors of messengers for intercellular communication (Bollini et al., 2018; Murphy et al., 2019; Tatischeff, 2019), the vacuolation and exocytosis in the regenerating blastema reminded us of the vesicle-mediated intercellular communication signals within the blastema. We partitioned the regenerating tissues into three subcellular fractionations, and analyzed the proteins and small RNAs components. Out of 1561 differential proteins (6dpa vs. 0dpa) identified from three subcellular fractions, 82 proteins were included in the 626 differential transcripts.

We then transferred each fraction into the in vitro cell cultures of the PAC2, and examined the cell transcription profile. We found that the induced transcription profiles were functionally comparable to the identified 82 proteins and 626 differential transcripts from the regenerating blastema, suggesting that soluble fractions (Fraction III and Fraction II) could repress Wnt, FoxO-autophagy signaling pathways, but enhanced cardiac muscle regeneration, MAPK and Hedgehog signaling pathways. Table S1 listed the equivalent functional images of the three core blastema genes and two histone marks. Significantly histone H3K9 demethylation genes *(kdm7aa, kdm7ab,phf8)* were induced by blastema subcellular fractions in the PAC2 cells.

Further functional clustering analysis of the 82 proteins showed the highest enriched ten extracellular molecules and five intermediate filaments (Table S6). Because some of these specific components were reported to accompany the cell shape change (thymoglobulin, lens crystallins), ECM, and cytoskeleton (keratin, vimentin) (Greenburg and Hay, 1988), and intermediate filament (IF) networks are implicated in both nuclear and cytoskeletal organization (Etienne-Manneville and Lammerding, 2017; Gruenbaum and Foisner, 2015), our data suggested that these extracellular components and Ifs may interact at focal adhesion complexes, and activate cytoplasmic-nuclear signal transduction pathways.

### 8. Krt5 targeted BMP4 signaling and regulated H3K9Ac/Me3 reciprocal chromatin deposition and core blastema gene activation

Since H3K9 methylation is an important epigenetic determinant for chromatin accessibility, and H3K9 methyltransferases are downstream targets of bone morphogenetic proteins (BMPs) (Chen et al., 2013), we directly tested the effects of Keratin 5 (Krt5) on regulation of BMPs signaling pathway and on H3K9Ac/Me3-marked chromatin architecture. Because FBS contains abundant BMPs, and noggin is an effective natural BMP antagonist (Chen et al., 2013), including BMP4, BMP6 and BMP9, we cultured PAC2 cells in FBS-contained culture medium with or without Krt5 and/or noggin. PAC2 cells did not show any Krt5 signals. In Krt5-treated PAC2 cells, the Krt5 showed typical coarse dot-like structures scattered through the cytoplasm, particularly displaying the string of beads-like structure surrounding the periphery of the nucleus (Fig.4A, B, Fig.S9). In contrast to the noggin treated responses, Krt5 increased F-actin assembly around the nuclei, increased the ratio of H3K9Me3-positive cells, and decreased the pSmad staining intensity. These results suggest that Krt5 promoted F-actin labeled cytoskeleton reorganization and H3K9Me3 depositions via a noggin-differential BMP signaling pathway.

**Figure 4.**
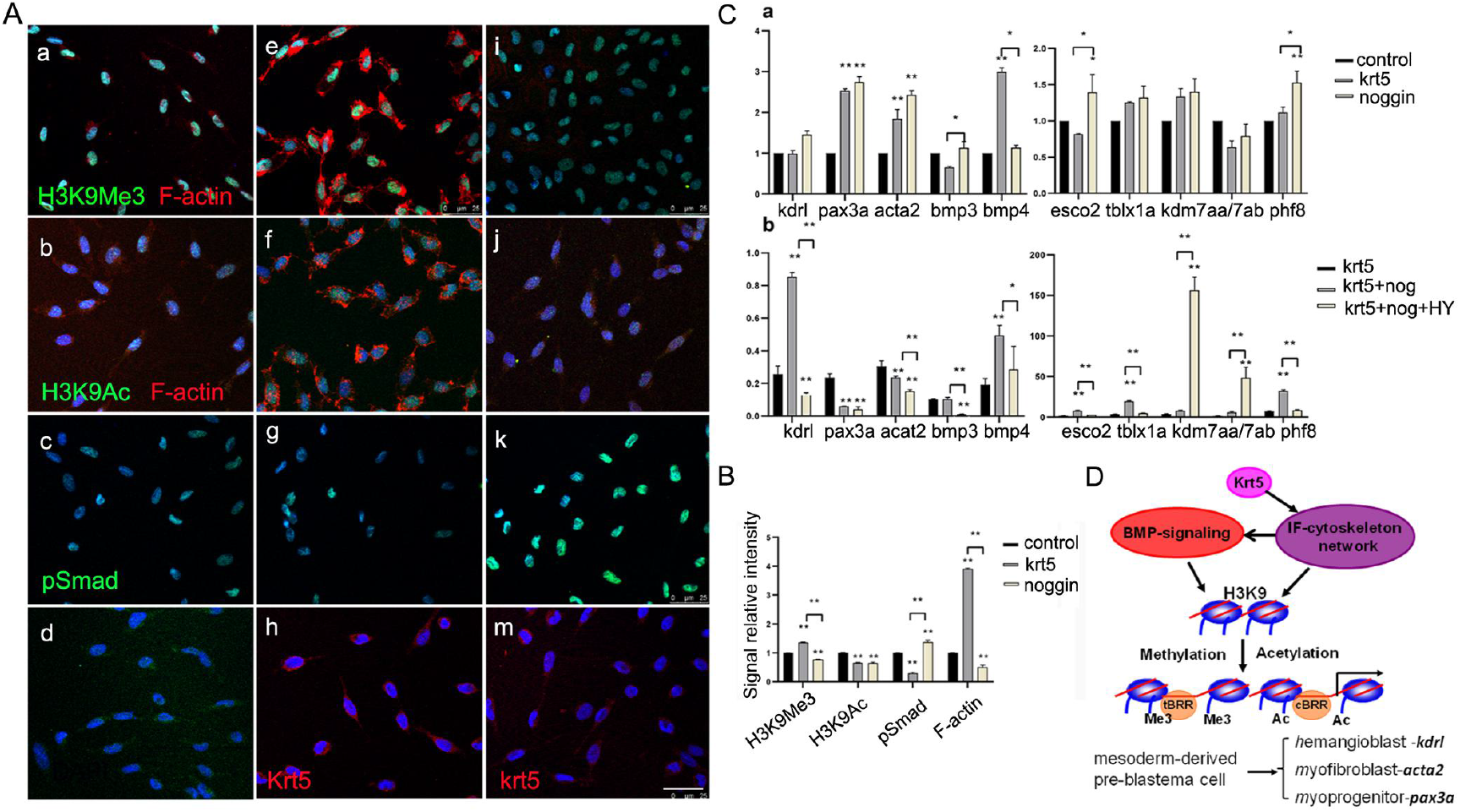
Effects of human Krt5, noggin and liothyronine on two histone marks and three core blastema genes in cultured PAC2 cells. A showed immunohistochemistry analyses. a-d, Control PAC2 cells cultured in 6%-FBS medium, e-h, PAC2 cells cultured in 6%-FBS medium with krt5, i-k, PAC2 cells cultured in 6%-FBS medium with noggin, m, PAC2 cells cultured in 6%-FBS medium with krt5 and noggin. DAPI staining blue. B showed quantification of A. C showed quantitative RT-PCR analyses. (a) PAC2 cells were separately cultured with Krt5, or noggin in 6%-FBS medium. The simultaneously cultured PAC2 cells without any treatment were taken as control. (b) PAC2 cells were cultured with Krt5 or cultured with Krt5 mixture with noggin and/or liothyronine in 12%-FBS medium. D. A proposed Keratin-BMP mechanosignaling to bridge extracellular-cytoplasmic signal transduction pathways and H3K9 acetylation/methylation chromatin switch in regulation of core blastema genes during blastema formation and tissue respecification. tBRR, temporal blastema respecification regulator; cBRR, constitutive blastema respecification regulator. Bar = 25 μm

We further examined the effects of Krt5-BMP signaling network on expression of three core blastema genes and two histone modification-related genes (Fig. 4C). Krt5 and noggin reversely regulated transcription of *bmp4, bmp4, kdrl*, *esco2* and *phf8.* When FBS content increased from 6% to 12%, these reverse effects were more significant on three core blastema genes and two histone modification-related genes. Furthermore, we showed that Krt5 itself had little effects on the expressions of the tested histone modification enzymes but enhanced noggin’s effects on histone acetyltransferase (*esco2*), deacetylase *(tblxr1a)*, and demethylase *(phf8, kdm7ab)* activities. Consistently Krt5 and noggin mixture increased *kdrl* and *bmp4* expression, and suppressed *pax3a* and *acta2* expression. As BMP signal transduction could be regulated by potent intracellular and extracellular antagonists (Yu et al., 2010), such differential regulatory effects between Krt5 and noggin confirmed that Krt5 did not take classic transmembrane receptor pathway to inhibit phosphorylation of Smad, but activated non-canonical BMP4 signaling pathway through intracellular mechanosignal transduction mechanisms in a dosage dependent manner.

## Discussion

Cell plasticity studies propose a desirable regenerative strategy which relies on the availability of stem cells and/or on the dedifferentiation and/or transdifferentiation of differentiated cells to replenish the missing structure (Galliot and Ghila, 2010). For heart regeneration after myocardial infarction, various anti-inflammatory agents, and stem cell and gene therapies have been tested to promote myocardial recovery and reduce reperfusion injury (Parviz et al., 2019), and yet no conclusively effective strategies have been established. In this study we found that zebrafish heart ventricle resection also encounters bleeding, hemostasis, and blood clot. Our findings indicate that zebrafish blastema regeneration could integrate inflammatory reaction and epithelial-mesenchymal transition (EMT) into cell remodeling network to salvage the damaged myocardium and resolve blood clot.

First, zebrafish can faithfully repair the complicated tissues and organs through regeneration blastema, a mass of pluripotent mesenchymal cells (Govindan and Iovine, 2015; Lepilina et al., 2006; Wang et al., 2012). There is much debate about the origins of the heart blastema cells, proliferation of the reserved undifferentiated cardiac progenitors, or transformation from differentiated myocardial and nonmyocardial cell types (Laflamme and Murry, 2011). The present study clearly showed that regeneration blastema arose from an advancing cell remodeling, which was marked by three representative blastema tissue specific factors (Flk1^+^hemoangiogenic progenitor, alpha-SMA^+^myofibroblast, and Pax3a^+^myogenic progenitor). During blastema formation alpha-SMA cells-dominant fibrogenesis provides a temporal tissue scaffold to pave the way for Flk1^+^ and Pax3a^+^ cell respecification within the blastema.

A significant observation on blastema cell remodeling was the coexistence of cytoplasmic efflux and nuclear reorganization in the damaged muscle cells. The extensive cytoplasmic degradation is accompanied by vacuole efflux and retention of the condense nucleus with small cytoplasm. Similar cytoplasmic vacuolation and efflux has once been reported in iris epithelial cell type conversion (Yamada, 1980). Pertaining to the blastema cytoplasmic lysis and exocytosis we also identified 28 proteolysis-related genes and lysosome-dependent mechanisms (Table S1, Fig.S10). Another significant observation was that the induced clot compartment showed very similar morphology to the previously described hematon, a multicellular functional unit of hematogenesis in normal mammalian bone marrow (Blazsek et al., 2000), which contains the putative hematopoietic stem cells and the stem cell niche (Chepko and Dickson, 2003).Supplementing this, several clusters of genes responsible for angiogenesis and hematopoiesis were significantly activated in both blastema tissue and induced fibroblast cells (Table S1).

Second, reverse histone modification patterns (such as methylation of H3K4 / H3K27, and H3K27Ac/ H3K27Me3) have been reported to regulate chromatin architecture and function in cell lineage specification (Hawkins et al., 2011; Li et al., 2008; Towbin et al., 2012). The present results suggested that reciprocal acetylation and methylation of H3K9 could regulate the blastema cell proliferation and respecfication through their specific enhancers and enhancer motifs (the key developmental and tissue specific regulators) (Fig.5C). Presumably those constant motifs were recruited to maintain the cell differentiation status while the temporal-sensitive motifs were responsible the blastema cell respecification. Consistent with the idea that H3K9 methylation is a barrier during somatic cell reprogramming into iPSCs, our data further suggested that H3K9Ac/Me3 switch delimit the blastema cell reprogramming at tissue-specific progenitors rather than pluripotent stem cells.

Finally, the present study can clarify why zebrafish blastema regeneration is epithelial-dependent (Wang et al., 2012; Zhang et al., 2015), and how undergo EMT (Lepilina et al., 2006). Our data showed that the epithelial-predominant keratin could help cell to bridge mechanosignaling between extracellular-cytoplasmic signal transductions and histone modifications-mediated chromatin remodeling, thus supporting the view that the keratin could modulate IF-cytoskeleton organization, interact with focal adhesions and cell-surface molecules (Quinlan et al., 2017; Sumer et al., 2019). By in vitro culture of zebrafish embryonic fibroblast cells, we demonstrated that the keratin 5 network could modulate H3K9Ac/Me3 chromatin state and expression of three core blastema tissue specific genes through the noncanonical BMP4 signaling pathway (Fig.5C). BMP signaling has long been acknowledged to be pivotal in mesoderm induction, vascular biology and hematopoietic commitment (Garcia de Vinuesa et al., 2016; Kirmizitas et al., 2017). Consistent with the reports that keratins could regulate yolk sac hematopoiesis and angiogenesis through BMP4 signaling (Vijayaraj et al., 2010), and BMP-Wnt signaling could specify hematopoietic fate through Cdx-Hox pathway (Lengerke et al., 2008), our result further suggest that keratin-BMP4-thyroid hormone cocktails induce mesoderm-derived embryonic fibroblasts undertaking blastema-like angiogenesis and musculature through dual regulation of cytoplasmic signaling and chromatin remodeling. Given that the blastema is a mixture of tissues, and PAC2 is a fibroblast cell line, our newly established keratin 5-BMP-H3K9Ac/Me3 mechanosignaling cascade should represent a basic mechanism of mesoderm-derived blastema regeneration, particularly toward blood system reconstruction. Our ongoing work will provide more detained mechanisms by identification of the intracellular Krt5-supercomplex.

Materials, methods, and any associated references are available in the supplemental data. All sequencing data have been deposited in the GEO repository (GSE98857) (https://www.ncbi.nlm.nih.gov/geo/query/acc.cgi?acc=GSE98857).

## Acknowledgements

We appreciate Professor Xin A. Zhang for editing the manuscript.We thank Kailun Lye and Xinyu Li for their technical assistances in ultrasctructural and subcellular fraction samples. This work was supported in part by Shanghai Universities First-class Disciplines Project of Fisheries, and National Natural Science Foundation of China (31772840, 31771431).

## Author contributions

Conceived and designed the experiments: JY. Performed the experiments: HG, XW, FY, YP, WZ. Analyzed the data: JY, HG, XW, FY. Wrote the paper: JY, HG, XW, GW, CW.

