## Supplementary material for "Keratin5-BMP4 mechanosignaling regulates reciprocal acetylation and methylation at H3K9 to define blastema cell remodeling during zebrafish heart regeneration": methods,supplemental figures and tables

### Materials and Methods

#### Animals

Zebrafish TAB lines, Flk1 promoter-derived green fluorescent protein (GFP) expression construct Flk1: GFP transgenic zebrafish (Flk1-transgenic fish) were reared as previously described. All experiments were performed in accordance with the Animal Care and Use Committee guidelines of Shanghai Ocean University (SHOU-DW-2016-004).

#### Sample process, histology and immunohistochemistry

After resection of the ventricular apex, the heart tissues were dissected at the day indicated. Histology sectioning, and staining for the immunohistochemistry (IHC) were performed as described previously (Zhang et al., 2015). For ultrastructural observations, heart tissues were dissected as mentioned above, and fixed 3 hours in Karnovsky's fixative at 4°C. After 6 PBS (phosphate buffer) washes, tissues were again fixed for 2 h in 1% osmium tetraoxide in phosphate buffer at 4°C. After several washes, the tissues were dehydrated in graded acetone solutions and embedded in Epoxy resin using Epoxy embedding medium kit (45359, Sigma, St. Louis, MO, USA). Ultra-thin sections of 60-80 nm thickness were stained with alcoholic uranyl acetate and lead citrate for appropriate time intervals. The grids were then examined with Transmission Electron Microscope HT7700.

#### Transcription analyses

Total RNA was isolated from tissues using TRIzol Reagent (Invitrogen) and further purified using RNeasy Mini Kit (Qiagen) and run on Agilent Bioanalyzer 2100 to asses sample integrity. The mRNA-seq library preparation from 1 μg of total RNA was performed with TruSeq RNA Sample Prep Kit (Illumina) according to the manufacturer's instructions, and 2x150 paired-end sequencing performed on Illumina Hiseq 2500. For RNA-Seq data Analysis, reads from RNA-seq experiments were aligned to zebrafish genome build danRer10 using

STAR (Dobin et al., 2013). Duplicate reads were removed with SAM tools (Li et al., 2009). The mRNA expression of a gene was quantified by FPKM (Fragments Per Kilobase of transcript per Million mapped reads) based on RefSeq gene annotation using Cuffdiff (Trapnell et al., 2013). Differential expressed genes were identified by a Fold change of >= 1.5 and FPKM of >= 3 for at least one of the two conditions based on R package edgeR. The GO enrichment and KEGG pathway analysis were performed using the R package cluster Profiler. The GO terms with FDR<0.05 were consider as significant. RT-PCR assays were performed as described (Zhang et al., 2015).

**Chromatin immunoprecipitation (ChIP) -sequencing**

Chromatin was prepared from fresh heart tissues following the manual instruction of ChIP-IT High Sensitive Kit (Cat.No. 53040, Active Motif. Tokyo, Japan). ChIP assays were carried out using previously described protocols (Wei et al., 2009). For each sample, fixed tissues were lysed to prepare nuclear extracts. After chromatin shearing by sonication, lysates were incubated overnight at 4°C with protein A Dynabeads (Invitrogen) coupled with 3-5 μg of antibody specific for H3K9Ac or H3K9Me3. Recovered DNA was analyzed by quantitative PCR (ChIP-qPCR) and sequencing (ChIP-Seq). ChIP-seq libraries were generated by ThruPLEX-FD Prep Kit (Product No.R40012, Rubicon, Ann Arbor, USA), and sequenced by Illumina Hiseq 2500 using the paired-end module and with 150 bp reads on each end. Sequence reads of 150bp were obtained, mapped to the zebrafish genome (danRer10), and processed as described previously.

All ChIP-Seq data sets were trimmed and aligned to zebrafish genome build danRer10 genome using Bowtie2 (Langmead and Salzberg, 2012). Duplicate reads were removed with SAM tools. The Model-based Analysis for ChIP Sequencing v2.0 (MACS2) (Zhang et al., 2008) was used to identify regions of ChIP-Seq enrichment over background in an unbiased manner. For histone modification H3K9Ac and H3K9Me3 ChIP-Seq, we modified the parameters to facilitate accurate detection of broad peaks (--broad --broad-cutoff 1E-3 -p 1E-5). The top 10000 peaks were used for known motif enrichment analysis, which were performed using HOMER (v4.9) (Heinz et al., 2010) with default parameters. Other analysis was performed using deep tools (Ramirez et al., 2014). To identify regions that were differentially enriched, the HOMER software was used (Heinz et al., 2010). Super enhancers (SEs) were identified using ROSE (https://bitbucket.org/young_computation/rose). Closely spaced peaks (except those within 2 kb of TSS) within a range of 12.5 kb were merged together, followed by the measurement of H3K9Ac signals. These merged peaks were ranked by H3K9Ac signal and then classified into SEs or (typical enhancers) TEs. Both SEs and TEs were assigned to the nearest refSeq genes.

**Partition of subcellular fraction of blastema tissues**

The ventricular apex of 60 adult fish were amputated and reared for six days. The newly regenerated heart tissues were removed into 200ul of cold PBS, and minced. The homogenate was serially centrifuged to collect large cell components at 2,000g for 15 minutes (Fraction II). The supernatant was further centrifuged at 16,000g for one hour to collect extrudants and EV pellets. For cell culture, the pellets were redissolved in 200ul PBS and homogenized. For biomolecule identification, the collected supernatant (fraction II) or pellets (fraction III) were separately dissolved in Trizol for RNA extraction and protein isolation according to the manufacturer's manual. The yield RNAs and proteins were used for transcriptome analyses and protein mass-spectrometer as previously described. The other 60 unampumtated fish were used as control.

**Cell culture**

Zebrafish embryonic fibroblast cellline (PAC2) were cultured in L-15 medium supplemented with 12% FBS at 32 C. To test the effects of the tissue blastema extract on PAC2 transcription, the collected subcellular fractions were separately added into the PAC2 cultures. After 36 hours, the treated cells were washed, and suspended in Trizol for RNA extraction and RNA sequencing as described above. To test Krt5-BMP signaling networks, Krt5, recombinant human noggin, Liothyronine and antibodies to Krt5 and phosphor-Smad1/5/8 were purchased. Krt5, noggin and liothyronine were used at concentrations of 300ng/ml, 200ng/ml, 200 µM, respectively.

**Image quantification and statistical analyses**

IHC-stained tissue sections were imaged on a confocal microscope (Leica TCS SP8). Signal intensity was quantified using ImageJ software. All experiments were carried out in triplicate and repeated at least twice. Statistical analyses were performed using SPSS 17.0 software. The antibodies and primers used are listed in Supplemental table 7.

**Supplemental figures and legends**

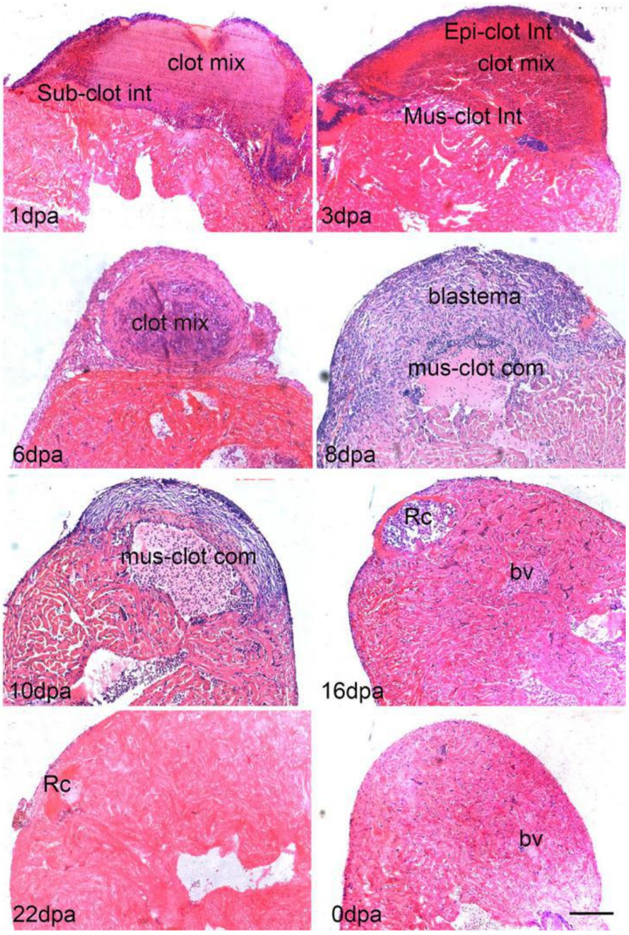


Fig. S1 Morphological analysis of heart regenerative process and blood clot replacement by Haematoxylin-Eosin. Bar = 200 μm.

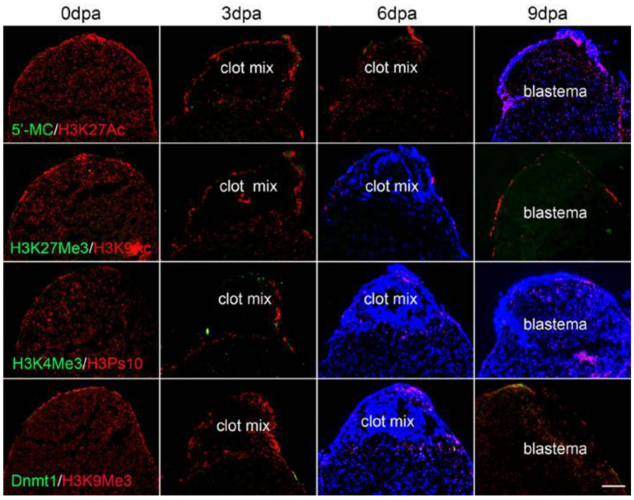


Fig. S2 Eight common epigenetic marks during heart regeneration using immunohistochemistry (IHC).

Tissue samples were collected at 0hpa, 3dpa, 6dpa and 9dpa. The immunostaining intensity of each epigenetic mark in normal uncut heart (0dpa) was used as mean basal levels. The specific antibodies for the following epigenetic marks were used: DNA methylation (5-methylcytosine, 5-MeC; cytosine-5-methyltransferase 1, DNMT1), histone acetylation (H3K9Ac, H3K27ac), histone methylation (H3K4Me3, H3K27Me3, H3K9Me3) and histone phosphorylation (H3PS10). Relative to temporal emergence of 5mC and H3K4Me3 at 3dpa in small number of light cells at intersections of epi-clot and mus-clot, all neotissue cells including the clotted cells showed stable H3K27Ac expression, and global loss of H3K27Me3 and DNMT1 during heart regeneration from 3dpa to 9dpa. H3PS10 expression was detected at all stages in part of the cells, presumably in the proliferating cells. Bar = 100 μm.

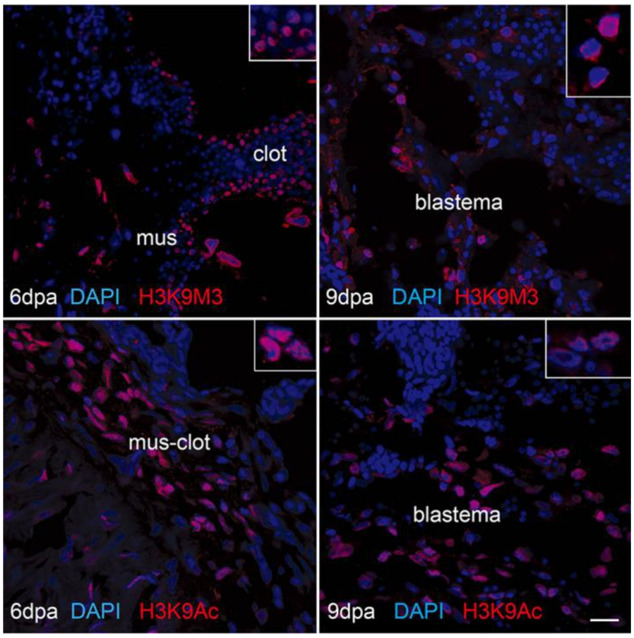


Fig. S3 Subcellular localizations of H3K9Ac and H3K9Me3 within the transforming cells. Inlet showed magnification. Bar = 25 µm.

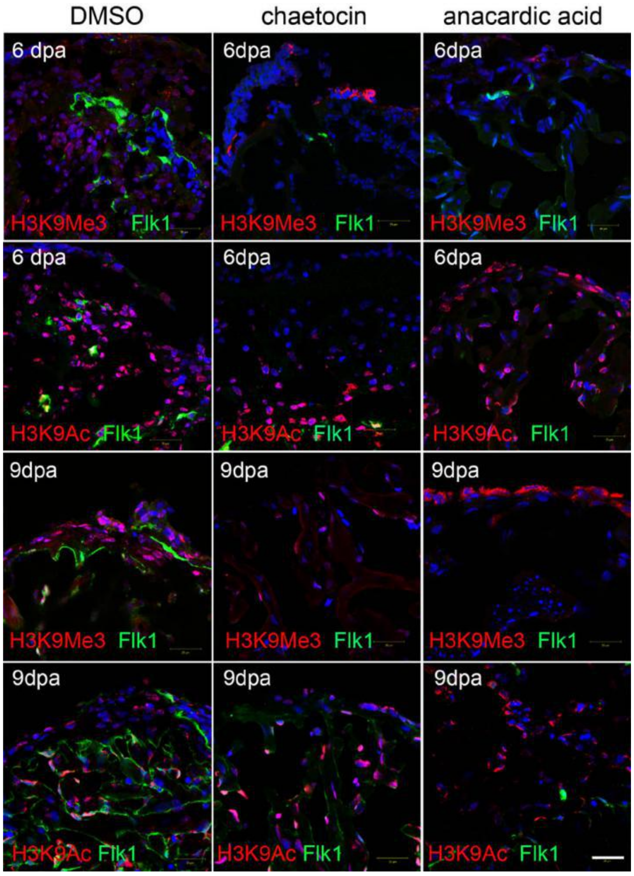


Fig. S4 Antagonist pretreatments altered H3K9Me3 and H3K9Ac depositions and Flk1-GFP distribution.

Chaetocin and anacardic acid were used as a specific inhibitors of the lysine-specific histone methyltransferase and histone acetyltransferase respectively. After ventricle amputation, four fish each group were soaked in DMSO (0.05%), chaetocin (500nM), or anacardic acid (500 nM), IHC was conducted at 6dpa. Bar = 50 μm

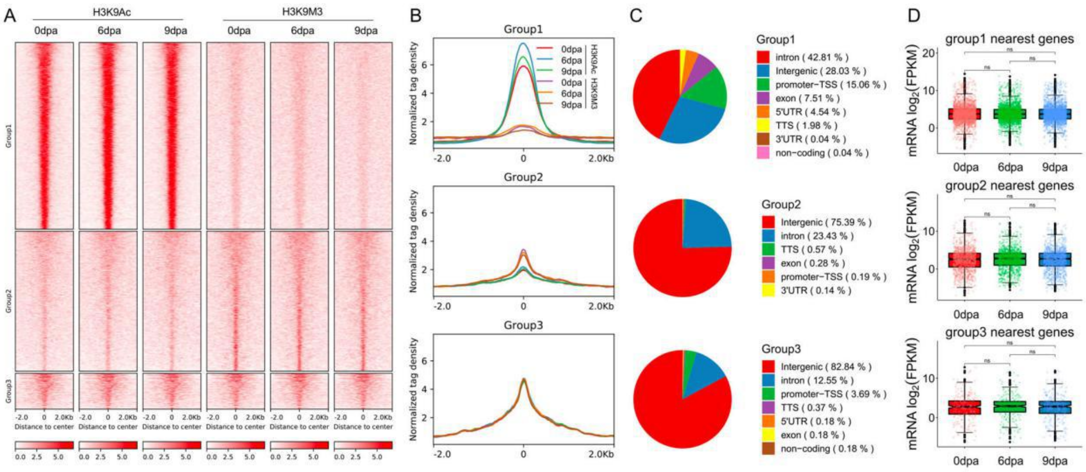


Fig. S5 ChIP-seq identification of H3K9Ac- and H3K9Me3-specific intergenic enhancers.

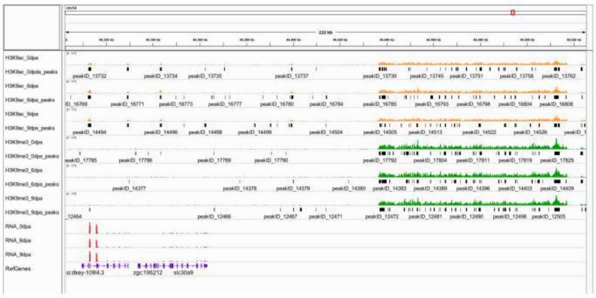

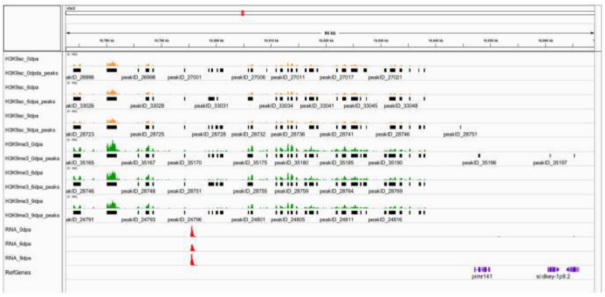

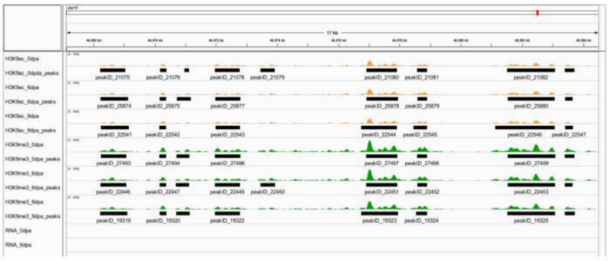

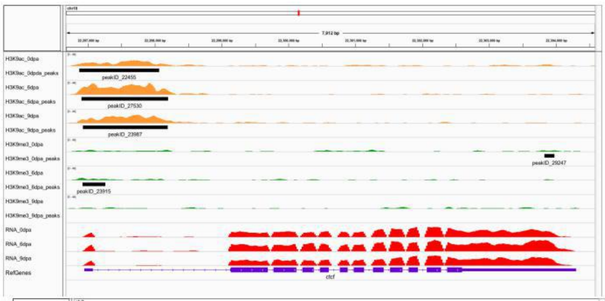

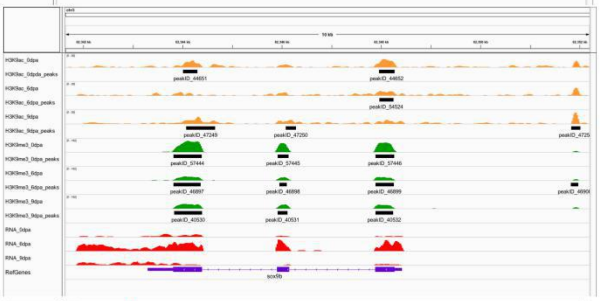

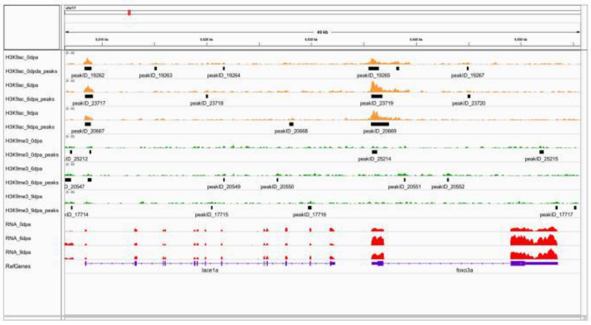

Fig. S6 Enrichments of H3K9Ac and H3K9Me3 marks at the nearest genes.

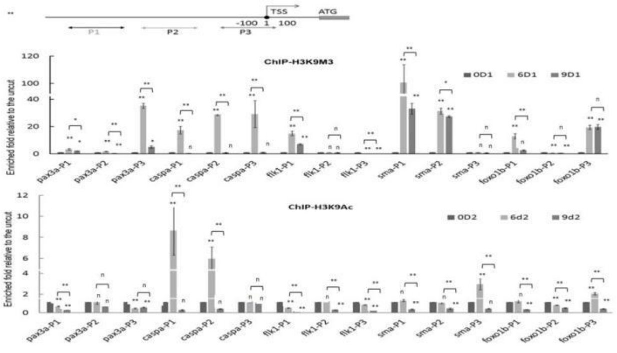


Fig. S7 ChIP-PCR reevaluation of the enrichment of the two histone modification marks at the promoters of the target genes. Flk1 (*kdrl*), alpha-SMA (*acta2*).

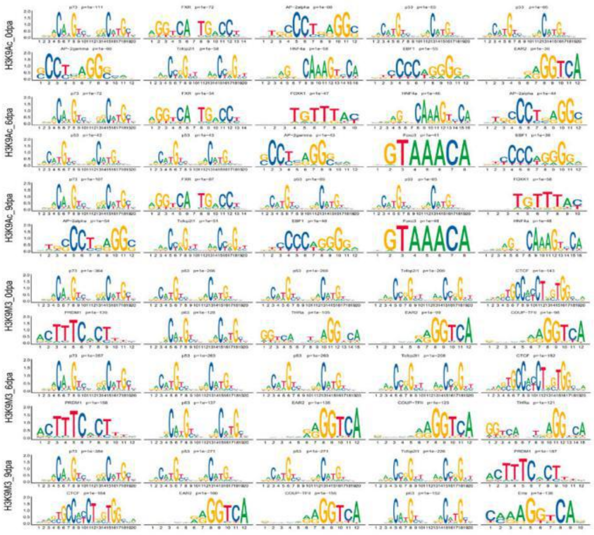

Fig. S8 Predictions of H3K9Ac- and H3K9Me3-specific motifs.

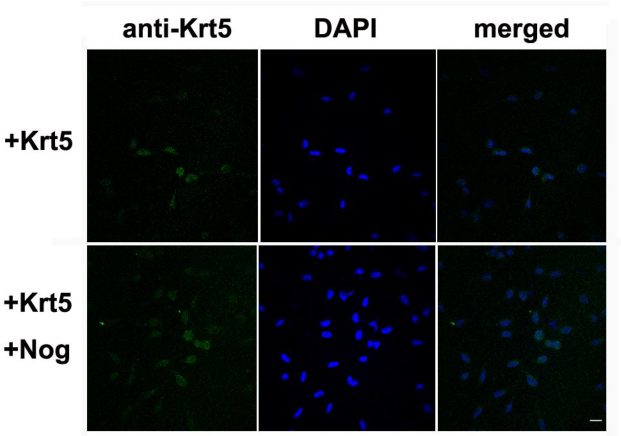


Fig. S9  Localization of the Krt5 peptide in the Krt5 and/or noggin transfetced PAC2 cells. Bar = 25 μm.

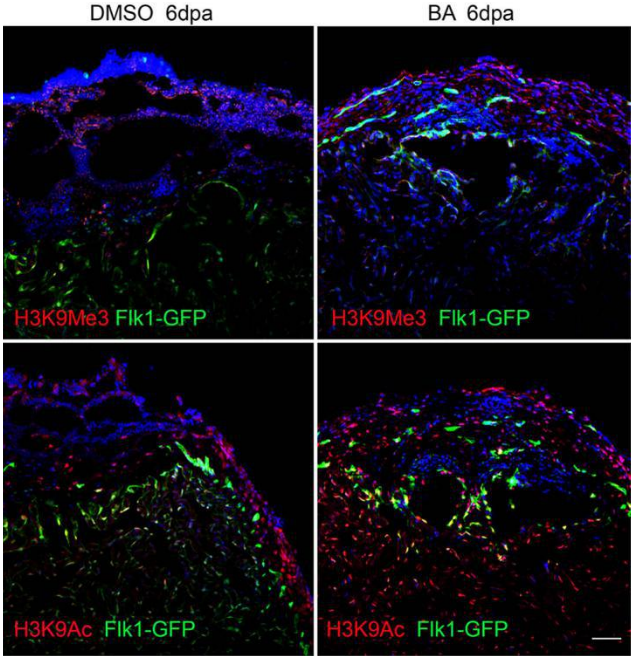


Fig. S10 Bafilomycin A1 pretreatment reduced tissue degradation, and enhanced Flk-1 indicated angiogenesis.

Bafilomycin A1 pretreatment of amputated heart reduced tissue degradation and exaggerated flk1-expressing vasculature. Bafilomycin A(1) is a specific inhibitor of the vacuolar type H(+)-ATPase (V-ATPase) in cells, and can inhibit the acidification of organelles containing this enzyme, such as lysosomes and endosomes(Klionsky et al., 2008). Bar = 50 μm

**Supplemental tables**

Table S1 Functional comparison of identified genes between blastema tissues and subcellular fractions-induced PAC2 cells.

Table S2 H3K9Ac and H3K9Me3 reciprocal depositions at chromatin.

Table S3 H3K9Ac and H3K9Me3 enrichments and the nearest genes' transcription.

Table S4 Corresponding motifs at the predicted H3K9Ac-/H3K9Me3-specific enhancers

Table S5 Prediction of H3K9Ac-/H3K9Me3-specific enhancer motifs at three core blastema genes

Table S6 Key blastema effectors identified by functional annotation clustering of the subcellular fraction proteins.

According to Human intermediate filament database, http://www.interfil.org/proteins.php, other Ifs, such as, vimentin (vim), desmin (desma), and synemin (synm) were identified in the partitioned cell components.

Table S7 Antibodies, primers and other agents used in this study.
